# Dominant nonsense mutations in efemp1 alter vertebral and craniofacial characteristics in adult zebrafish

**DOI:** 10.1101/2024.02.16.580611

**Authors:** Arianna Ericka Gómez, Kurtis Alvarado, Rohda Ahmed Yase, Yi-Hsiang Hsu, John D. Hulleman, Ronald Young Kwon

**Author notes:** Corresponding author, Ronald Young Kwon, PhD, 850 Republican Street, Box 358056, Seattle, WA 98109.

## Abstract

Heritable Disorders of Connective Tissues (HDCT) are a heterogenous, pleiotropic group of conditions that broadly affect connective tissues. *EFEMP1* is a member of the fibulin family of extracellular matrix (ECM) glycoproteins which is expressed in various human tissues. Individuals with *EFEMP1* variants have recently been identified and appear to have Marfan-like characteristics. Clinical phenotypes of these individuals include hernias, advanced bone age, tall stature, myopia, joint laxity, and thin skin. *EFEMP1*-associated HDCTs have been identified in individuals with biallelic and monoallelic variants. There is an urgent need to better understand the role of *EFEMP1* in regulating connective tissues including bone, and the pathophysiological mechanisms underlying the spectrum of genotype-phenotype relationships seen in EFEMP1-associated HDCTs. To investigate the role of *EFEMP1* in developing and adult bone, we used CRISPR-based editing to generate two efemp1 zebrafish alleles encoding for premature termination codons (PTCs) predicted to delete or severely alter the fibulin-type domain. Both alleles exhibited similar phenotypes in juvenile and adult fish. In juvenile fish, we did not identify changes in body size or vertebral development. In adults, we found significant changes in body length, bone microarchitecture, and craniofacial measurements in both heterozygous and homozygous mutant fish. These results expand our understanding of the role of *efemp1* in the skeleton and highlight the potential for dominant nonsense variants to play a role in manifestation of clinical phenotypes in *EFEMP1*-associated HDCTs.

## Introduction

Heritable Disorders of Connective Tissues (HDCT) are a heterogenous, pleiotropic group of conditions that broadly affect connective tissues (National Academies of Sciences et al. 2022). EGF-containing fibulin-like extracellular matrix protein 1 (EFEMP1) is an extracellular matrix (ECM) glycoprotein encoded by the *EFEMP1* gene. Also known as fibulin-3, EFEMP1 is a member of the fibulin family of proteins, which are characterized by tandem EGF domains and a C-terminal fibulin module (Timpl et al. 2003). Recently, biallelic and recessive nonsense and missense variants in *EFEMP1* have been identified in four individuals with Marfan-like clinical phenotypes (Mégarbané et al. 2012; Bizzari et al. 2020; Driver et al. 2020; Verlee et al. 2021). Phenotypic features in affected individuals include craniofacial characteristics, joint laxity, kyphoscoliosis, arachnodactyly, hyperextensible skin, inguinal hernia, and tall stature (Forghani et al.). These features are reminiscent of Marfan syndrome, an autosomal dominant condition. The majority of individuals with Marfan syndrome carry missense or nonsense variants in fibrilin-1 (*FBN1*). While several clinical phenotypes overlap in Marfan syndrome and individuals with *EFEMP1* variants, the EFEMP1-associated condition does not have a cardiovascular phenotype which is typically observed in individuals with Marfan syndrome. Conversely, Marfan syndrome is not associated with inguinal hernias which are observed in individuals with the EFEMP1-associated condition. Thus, *EFEMP1*-associated HDCTs are a newly discovered class of conditions that are both genetically and phenotypically distinct from other HDCTs.

The spectrum of variants underlying *EFEMP1*-associated HDCTs is rapidly expanding. Until recently, *EFEMP1*-associated HDCTs had been identified in four individuals with homozygous or biallelic recessive variants. However, Forghani et al. recently identified a heterozygous autosomal dominant-form of an *EFEMP1*-associated HDCT (Forghani et al.). Specifically, a proband and her mother with comparable phenotypic features were both heterozygous for a nonsense variant (p.Arg362*). While recent *in vitro* studies have suggested that the homozygous recessive C55R variant observed by Bizzari et al causes aberrant disulfide bonding in EFEMP1 (Woodard et al. 2022), how variants with different modes of inheritance may underlie *EFEMP1*-asssociated HDCTs are only partially explained by animal models. For instance, in the murine *Efemp1^-/-^* model, phenotypes included inguinal hernias, reduced elastic fibers in connective tissue, and accelerated aging phenotypes (McLaughlin et al. 2007). The similar phenotypes in *Efemp1^-/-^* mice and individuals with *EFEMP1*-associated HDCTs suggests that biallelic loss-of-function in *EFEMP1* is sufficient to cause the Marfan-like condition. However, mice heterozygous for the *Efemp1* deletion did not overtly develop hernias observed in homozygous mutants, arguing against *EFEMP1* haploinsufficiency. Animal models demonstrating the potential for heterozygous nonsense variants in *EFEMP1* to give rise to the Marfan-like condition are currently lacking.

Based on the characteristics of humans carrying *EFEMP1* variants, i.e. tall stature, lengthened bones, advanced bone age, and craniofacial characteristics, it is conceivable that some clinical features observed in EFEMP1-associated HDCTs involve altered function in bone formation or turnover. ECM proteins play a role in mediating signals to osteoclasts, osteoblasts, and osteocytes, cells that regulate bone remodeling (Alford et al. 2015). Perturbation of ECM glycoproteins in animal models results in abnormal bone remodeling and bone microarchitecture (Delany et al. 2000; Hankenson et al. 2000; Amend et al. 2015). In mice, *Efemp1* is expressed in developing bone and cartilage, and has been found to negatively regulate chondrocyte differentiation in a carcinoma-derived chondrogenic line (Ehlermann et al. 2003; Wakabayashi et al. 2010). However, evidence from knockout animal models pointing to a role for EFEMP1 in regulating bone in animal models remains equivocal. For instance, when on the BALB/c background, *Efemp1^-/-^* mice displayed curved spines with advanced age and reduced bone density (McLaughlin et al. 2007). However, these phenotypes were not observed in *Efemp1^-/-^* mice in the C57BL/6 background (McLaughlin et al. 2007). Moreover, altered craniofacial morphology in *Efemp1^-/-^* mice was not reported. Thus, there is limited evidence that loss-of-function in *EFEMP1* contributes to the skeletal and craniofacial characteristics observed in individuals with *EFEMP1*-associated HDCTs.

Here, we assessed the effects of *efemp1* mutations in zebrafish, a teleost whose small size and genetic tractability are amenable to rapid-throughput studies of the juvenile and adult skeleton. We hypothesized that heterozygous and homozygous nonsense mutations in *efemp1* would alter vertebral and craniofacial phenotypes. We isolated *efemp1* alleles encoding for PTCs and determined skeletal changes in juvenile and adult animals. We demonstrate altered vertebral and craniofacial characteristics in adult zebrafish for two different nonsense mutations in *efemp1*. Importantly, phenotypic severity in animals with heterozygous mutations in *efemp1* is comparable to homozygous mutants. Our studies support the potential for loss-of-function in *EFEMP1* to contribute to altered skeletal and craniofacial characteristics in individuals with *EFEMP1*-associated HDCTs, and suggest that heterozygous nonsense variants could play a role in the manifestation of clinical phenotypes.

## Materials and Methods

### Ethics Statement

Approval for this project was granted by the University of Washington Institutional Animal Care and Use Committee (IACUC) under protocol #4306-01.

### Animals

Zebrafish were housed on a 14:10 hour light:dark photoperiod at a 28.5°C temperature. Fish were housed in plastic tanks on a commercial recirculating aquaculture system and fed a commercial diet. All experiments were performed in mixed sex wildtype (AB) and mutant (*efemp1^+/w1014^* and *efemp1^+/w1016^*) lines on an AB background.

### CRISPR-based gene editing

To generate *efemp1* mutant alleles, we performed CRISPR-Cas9 gene editing as previously described (Watson et al. 2020; Watson et al. 2022). We used a guide RNA (gRNA, Alt-R system, Integrated DNA Technologies) targeting exon 4 of *efemp1* complexed with Cas9 protein (HiFi 3xNLS-Cas9, IDT). We mixed the gRNA with the crRNA and tracrRNA in a 1:1 ratio, followed by a 5-minute incubation at 95°C for 5 minutes, and cooling on ice. The ribonucleorotein (RNP) complex was made by combining the crRNA:tracrRNA:gRNA mixture in a 1:1 molar ratio with the Alt-R S.p. Cas9 nuclease (IDT) containing a 3XNLS sequence and incubating for 5-10 minutes at room temperature. The Cas9:gRNA RNP complex had a final concentration of ∼25 μM for injection. The RNP complex mixture was loaded into pre-pulled microcapillary needles (Tritech Research). Following calibration, 2nL of RNP complex was injected into the yolk of 1- to 4-cell stage embryos. The *efemp1* gRNA used was 5’-CGTGTCTGGAGCCGTATGTG-3’.

### Isolation of mutant alleles

We identified somatic founders carrying mutated *efemp1*, which were used to generate F1 progeny. Sanger sequencing was performed in F1 progeny and two predicted loss of function alleles were identified: *efemp1^w1014^*(ENSDART00000082142.6: c.437_444del, (p.Val147Leufs*60)), *efemp1^w1016^* (ENSDART00000082142.6:c.434_437del, (p.Try146Cysfs*28)). Founders with identical alleles were inbred to create stable F2 germline *efemp1* mutants. Germline *efemp1*^w1014^ and *efemp1*^w1016^ lines were maintained through heterozygous inbreeding and clutches of mixed genotypes were housed together. Genotyping was performed using standard PCR conditions (35 cycles, 58°C annealing temperature) with the following *efemp1* primers: F: 5’-GGAGGAAACCCACACCAACA-3’, R: 5’-CAAACAGCGGATTGAGCGAC-3’. Gel electrophoresis was performed on a 2% agarose gel to resolve different sizes of PCR amplicons of wildtype, heterozygous, and mutant alleles.

### Vertebral mineralization analysis

Calcein labeling and imaging was performed as described previously (Watson et al. 2022). 13dpf larvae were stained with a 0.2% calcein (Sigma) solution in fish water for 20 minutes followed by 3x 10-minute washes in fish water. After staining, larvae were anesthetized in 0.01% MS-222 (Sigma) and mounted in borosilicate glass capillaries using 0.75% low-melt agarose (Bio-Rad) diluted in system water with 0.01% MS-222. We placed the capillary on a custom 3D printed holder for rapid orientation of the larvae. A high-content fluorescence microscopy system (Zeiss Axio Imager M2) and a 2.5x objective (EC Plan-Neoflaur 2.5x/0075) were used to take images. For each fish, a composite image stack was generated: 3/2 images in the x/y directions and optimized to ∼45 µm slice intervals in the z-direction across the entire region of interest, 13 slices all at 2.58 µm/pixel. Maximum intensity projections were generated from image stacks in Fiji for analysis. Fish were collected for genomic DNA extraction and genotyping after imaging.

Analysis was performed using Fiji. The polygon feature was used to mark the mineralized vertebral region and the area within this region was used to determine the area. Pixel units were converted to microns (1 pixel = 2.50 microns). In animals without mineralized vertebra, 0 microns was input as the measurement for that fish.

### RT-PCR

*efemp1* expression was assessed using the following primers: F: 5’-CCGGCTCCTACTACTGTGAG-3’, R: 5’-GCACATGTAGCTGGAGAACG-3’. For tissue specific expression, we isolated total RNA from AB wildtype fish which were dissected to collect bone, muscle, intestine, brain, skin, swim bladder, heart, eyes, and testes as described in (Gupta and Mullins 2010). For transcript expression of mutant alleles, we collected total RNA from whole adult AB wildtype, heterozygous, and homozygous mutant fish. RNA was extracted using Trizol-chloroform extraction, where tissues were homogenized in Trizol. cDNA synthesis was performed using the Superscript IV First Strand Synthesis System (Invitrogen). 1µL of cDNA was used for PCR using the DreamTaq Green DNA polymerase (ThermoFisher Scientific) in a 20µL reaction. Samples were run on a 2% agarose gel.

### microCT scanning and analysis

AB wildtype, heterozygous, and homozygous mutant animals were collected for microCT imaging at 90 dpf (Hur et al. 2017; Hur et al. 2018). We used the Scanco vivaCT 40 microCT scanner, with scan settings as follows: 21 µm voxel size, 55kVp, 145mA, 1024 samples, 500proj/180°, 200 ms integration time. Four fish were scanned simultaneously in each acquisition. DICOM files were generated for individual fish and FishCuT analysis was performed as previously described (Hur et al. 2017; Hur et al. 2018). Maximum intensity projections were used to measure standard length.

### Craniofacial measurements

Craniofacial shape variation was performed using a landmarking method we have previously described (Diamond et al. 2022; Diamond et al. 2023). We selected 14 landmark points to perform our analysis, which were derived from (Diamond et al. 2022). These 14 landmark points were manually placed on craniofacial landmarks using the markups module in 3DSlicer. For each fish, these landmarks were used to measure distance between specific points listed in Table 1 and 2. These measurements were selected based on prior work in zebrafish mutants (Diamond et al. 2022; Diamond et al. 2023). In some cases, measures were normalized for each fish by dividing by standard length.

**Table 1.** Location of individual landmarks used for craniofacial measurements. The 13 landmarks located on the zebrafish cranium used to measure differences craniofacial lengths in WT, HET, and MUT *efemp1^w1014^* and *efemp1^w1016^*fish.

| Landmark | Location |
| --- | --- |
| 1 | Anterior frontal bone |
| 2 | Dorsal frontal/parietal suture |
| 3 | Dorsal portion of first vertebrae |
| 4 | Left epiphyseal bar |
| 5 | Left posterior pterotic |
| 6 | Left opercle/interopercle suture |
| 7 | Left posterior opercle |
| 8 | Left anguloarticular |
| 9 | Right epiphyseal bar |
| 10 | Right posterior pterotic |
| 11 | Right opercle/interopercle suture |
| 12 | Right posterior opercle |
| 13 | Right anguloarticular |

**Table 2.** Reference points for craniofacial measurements. The landmarks used to measure specific distances on the zebrafish cranium. Landmarks can be visualized on a 3D rendering of a wildtype in Figure 4A.

| Landmark | Measurement |
| --- | --- |
| 1 – 3 | Skull length |
| 1 – 2 | Frontal/parietal bone length |
| 4 – 9 | Epiphyseal bar length |
| 8 – 13 | Distance between anguloarticulars |
| 7 – 12 | Posterior midline skull width |
| 5 – 10 | Dorsal skull width |
| 6 – 11 | Ventral skull width |

### Statistical approach

All statistical analyses were performed using GraphPad Prism. For each characteristic, a 2-way ANOVA was performed to determine p-values for genotype, allele, and genotype:allele interactions. In measurements where the genotype p-value was significant, we performed Tukey’s multiple comparisons test to perform pairwise comparisons.

## Results

### Isolation of *efemp1* mutant alleles

We first sought to define the expression of *efemp1* in adult tissues. We isolated 9 tissues from wildtype adult fish and performed RT-PCR, which identified *efemp1* expression in bone, intestine, skin, swim bladder, heart, eye, and testes (Fig 1A). The broad expression of *efemp1* suggested roles in multiple adult tissues including in bone, and motivated further studies to examine its function in mutant models.

**Figure 1.**
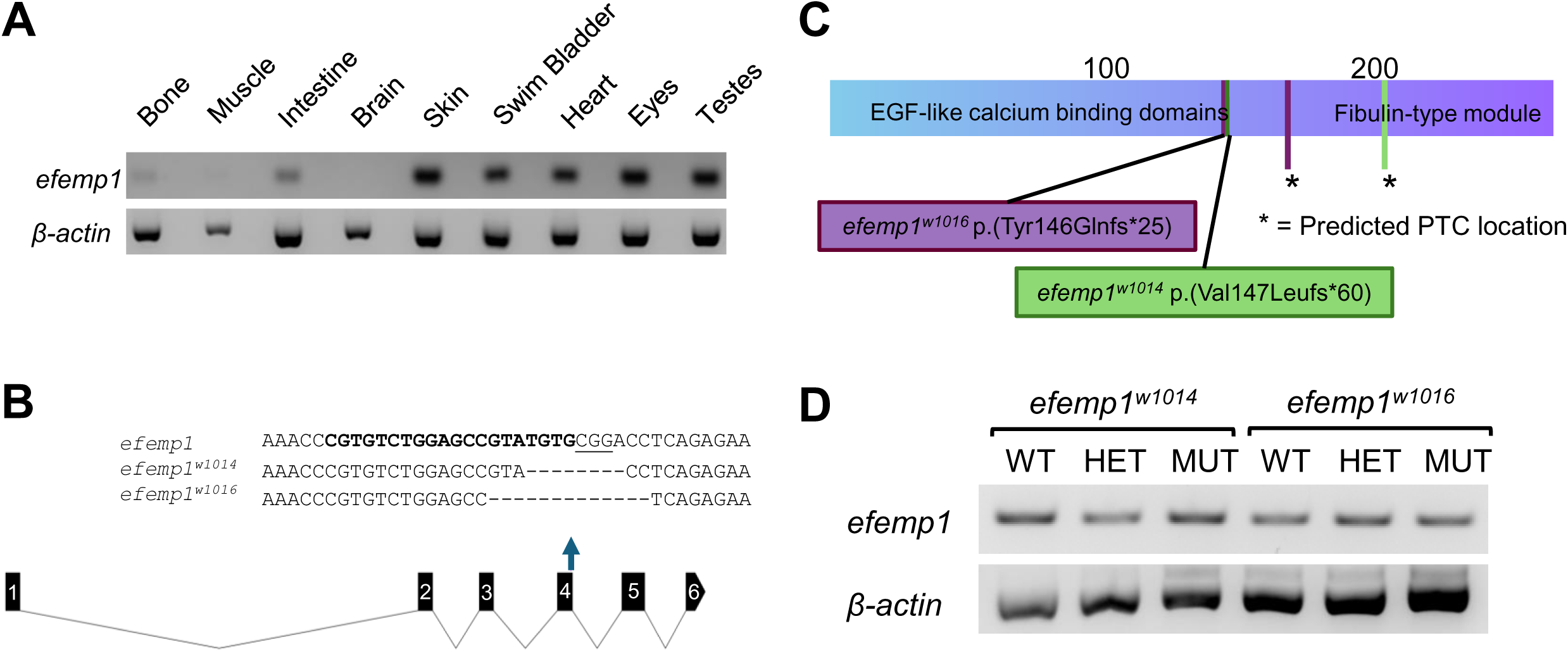
Expression of *efemp1* in adult zebrafish tissues and isolation of *efemp1* mutations in zebrafish. (A) RNA was isolated from adult zebrafish tissues to generate cDNA. Gel electrophoresis was performed using the cDNA to determine tissue specific expression. (A) Sequence of *efemp1* gRNA and the *efemp1^w1014^* and *efemp1^w1016^* alleles in the *efemp1* zebrafish gene. In the wildtype sequence, the bold bases indicate the gRNA sequence, and the underlined bases indicate the PAM sequence. (B) Mapped location of the *efemp1^w1014^* and *efemp1^w1016^* amino acid mutations and the downstream predicted termination codon (PTC) in the Efemp1 protein. (C) RT-PCR of wildtype (WT), heterozygous (HET), and homozygous mutant (MUT) *efemp1^w1014^* and *efemp1^w1016^* fish.

To generate mutant alleles, we designed an *efemp1* guideRNA (gRNA) targeting exon 4. Sanger sequencing in the F1 generation identified 2 *efemp1* mutant alleles with deletions in exon 4: *efemp1^w1014^*(c.437_444del; p.(Val147Leufs*60)) and *efemp1^w1016^* (c.433_445del; p.(Tyr146Glnfs*25)) (Fig 1B). Both mutations are predicted to cause a downstream premature termination codon in the protein sequence resulting in disruption or loss of the fibulin-type module (Fig 1C). Incross breeding of heterozygous fish carrying the *w1014* and *w1016* alleles produced offspring that were viable and were born with a Mendelian ratio.

To determine whether the mutated alleles were expressed in adult tissues, we performed RT-PCR in wildtype (WT), heterozygous (HET), and homozygous mutant (MUT) *efemp1^w1014^* and *efemp1^w1016^*fish. The *efemp1* transcript was expressed in fish carrying both alleles and in heterozygous and homozygous mutant fish, which suggests that the transcript is being transcribed in adult fish (Fig 1D). Thus, both alleles likely encode for a truncated protein product where the fibulin-type module is missing or severely disrupted. As the fibulin-type module is a conserved domain within the fibulin family, it is reasonable to assume that both alleles are functioning as hypomorphic or null alleles. Importantly, for all measurements reported below, no significant genotype:allele interaction, suggesting that the effects of *efemp1* mutations for both alleles were similar (Supp Table 1).

### Adult zebrafish with mutated Efemp1 exhibit larger body size

In 13 dpf juvenile fish, standard length was not significantly different between wildtype and mutant fish (Fig 2A). In adult 90 dpf *efemp1* mutants, there were significant differences in length between wildtype and mutant fish, where both were longer than control fish (Fig 2B and 2C).

**Figure 2.**
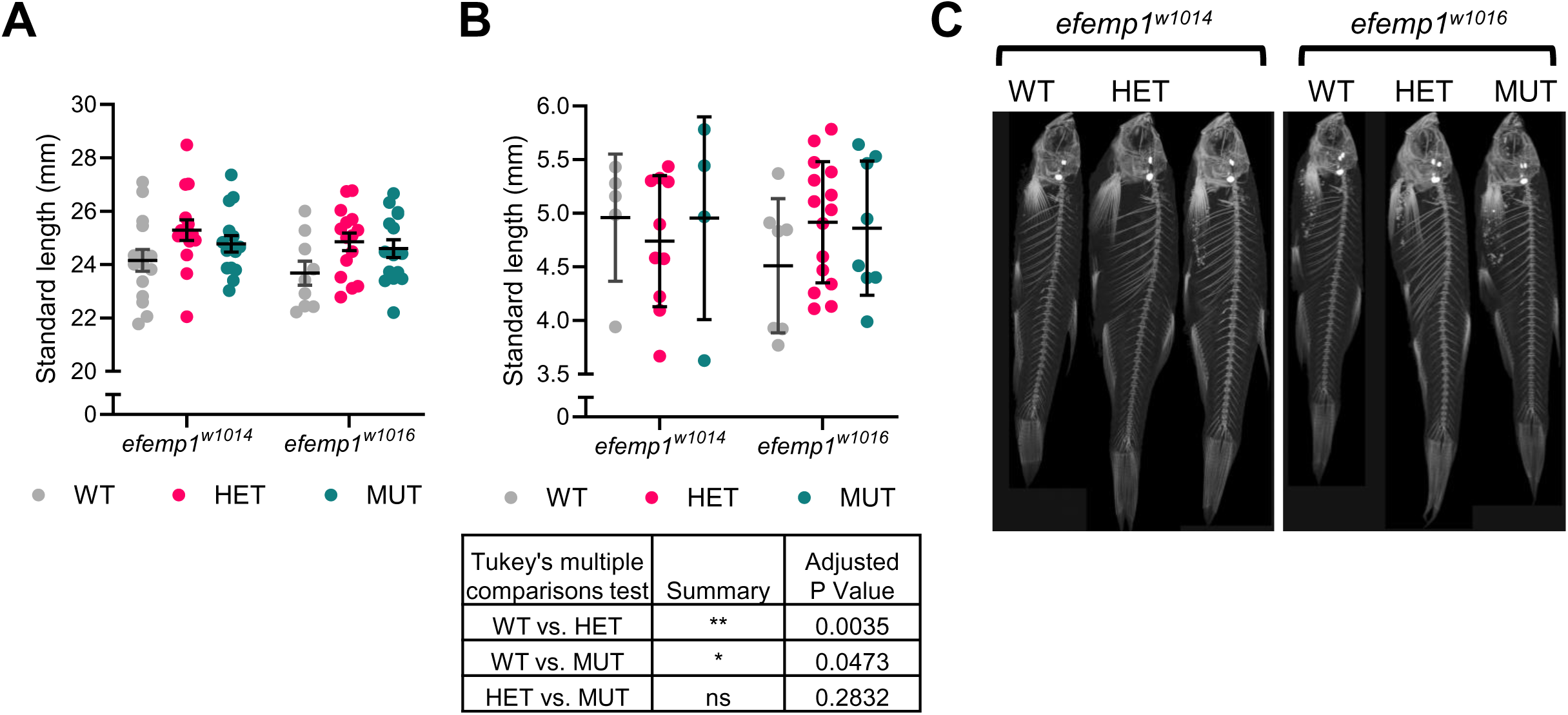
Standard length of juvenile and adult zebrafish carrying *efemp1* mutations. Standard length was measured in (A) 13 dpf and (B) 90 dpf WT, HET, and MUT *efemp1^w1014^* and *efemp1^w1016^* fish. In 90 dpf fish, a two-way ANOVA indicated significant differences between genotypes. The results of a Tukey’s multiple comparison test are shown below the graph. Each dot represents the standard length of a single fish. In (A) n = 4-15 fish per group and (B) n = 9-15 fish per group. (C) Representative max intensity projections of *efemp1^w1014^* and *efemp1^w1016^* microCT scans.

### Adult zebrafish with mutated Efemp1 are associated with altered vertebral morphology

In 13 dpf juvenile fish, we performed calcein staining, which stains mineralizing structures, and measured the area of mineralized vertebrae. We did not measure a significant difference in mineralized vertebra between wildtype and mutant fish (Fig 3A).

**Figure 3.**
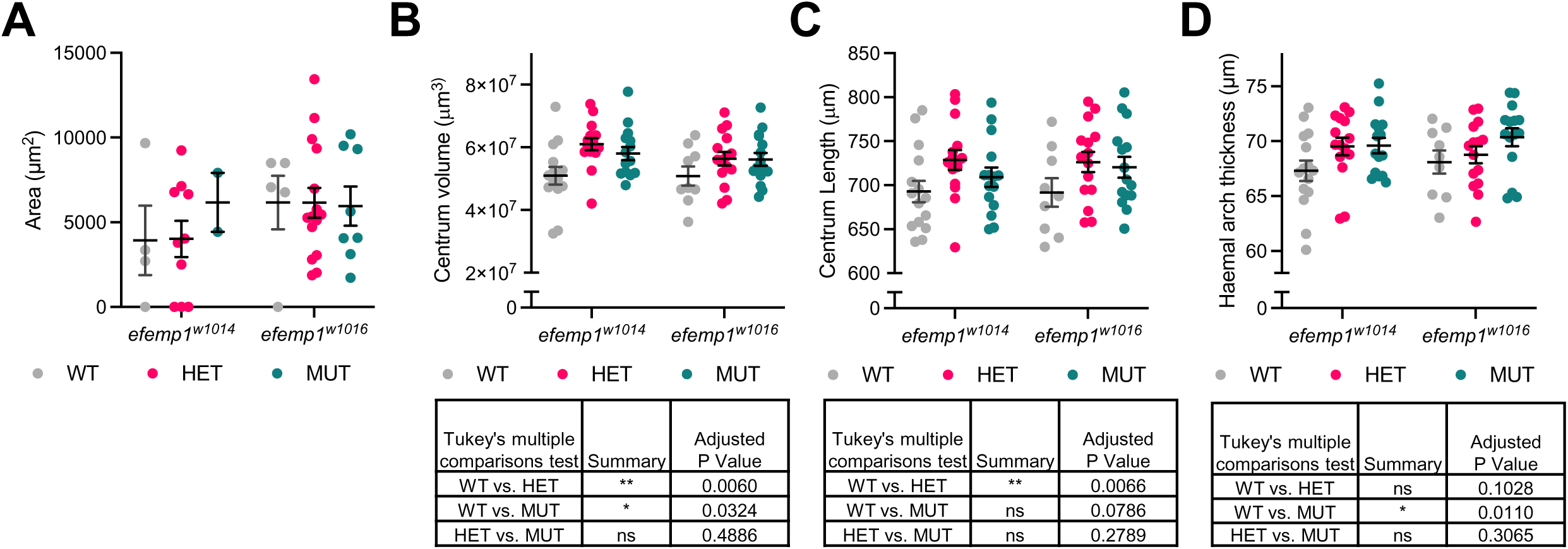
Vertebral characteristics of adult zebrafish carrying *efemp1* mutations. (A) Area of calcein staining in 13 dpf developing vertebra of *efemp1^w1014^* and *efemp1^w1016^* fish. Each data point is the average measured area of vertebrae for a single fish. n = 2-15 fish per group. FishCuT analysis results for (B) centrum volume, (C) centrum length, and (D) haemal arch thickness in *efemp1^w1014^* and *efemp1^w1016^* fish. The results of Tukey’s multiple comparisons test are shown below the graphs. Error bars represent mean and standard error of the mean. Data points represents the average measurement of a single fish for 16 vertebra. (B-D) n = 9-15 fish per group.

Since we identified expression of *efemp1* in adult bone, we were interested in the potential effects of mutated *efemp1* in the adult vertebral spine. In 90 dpf adult fish, we performed microCT scans and FishCuT analysis to determine changes in vertebral microarchitecture. We identified significant differences in centrum volume and centrum length in heterozygous mutants compared to wildtype fish (Fig 3B and 3C). Additionally, centrum volume and haemal arch thickness were significantly different when comparing wildtype and homozygous mutant fish (Fig 3B and D). Other vertebral microarchitecture measurements did not show significant differences between wildtype and mutant fish (Supp Fig 1). We did not observe an obvious difference in dysmorphic vertebrae in mutant fish compared to controls.

### Adult zebrafish with mutated Efemp1 are associated with altered craniofacial morphology

Utilizing microCT scans and methods previously developed by (Diamond et al. 2022), we measured the distance between specific craniofacial landmarks in wildtype and mutant fish. We measured the distance between 7 pairs of craniofacial landmarks and found significant changes in skull length and dorsal skull width in heterozygous mutant fish. Interestingly, *efemp1* mutations did not significantly affect homozygous mutant fish (Fig 4B). Other craniofacial measurements were not significantly different (Supp Fig 2).

**Figure 4.**
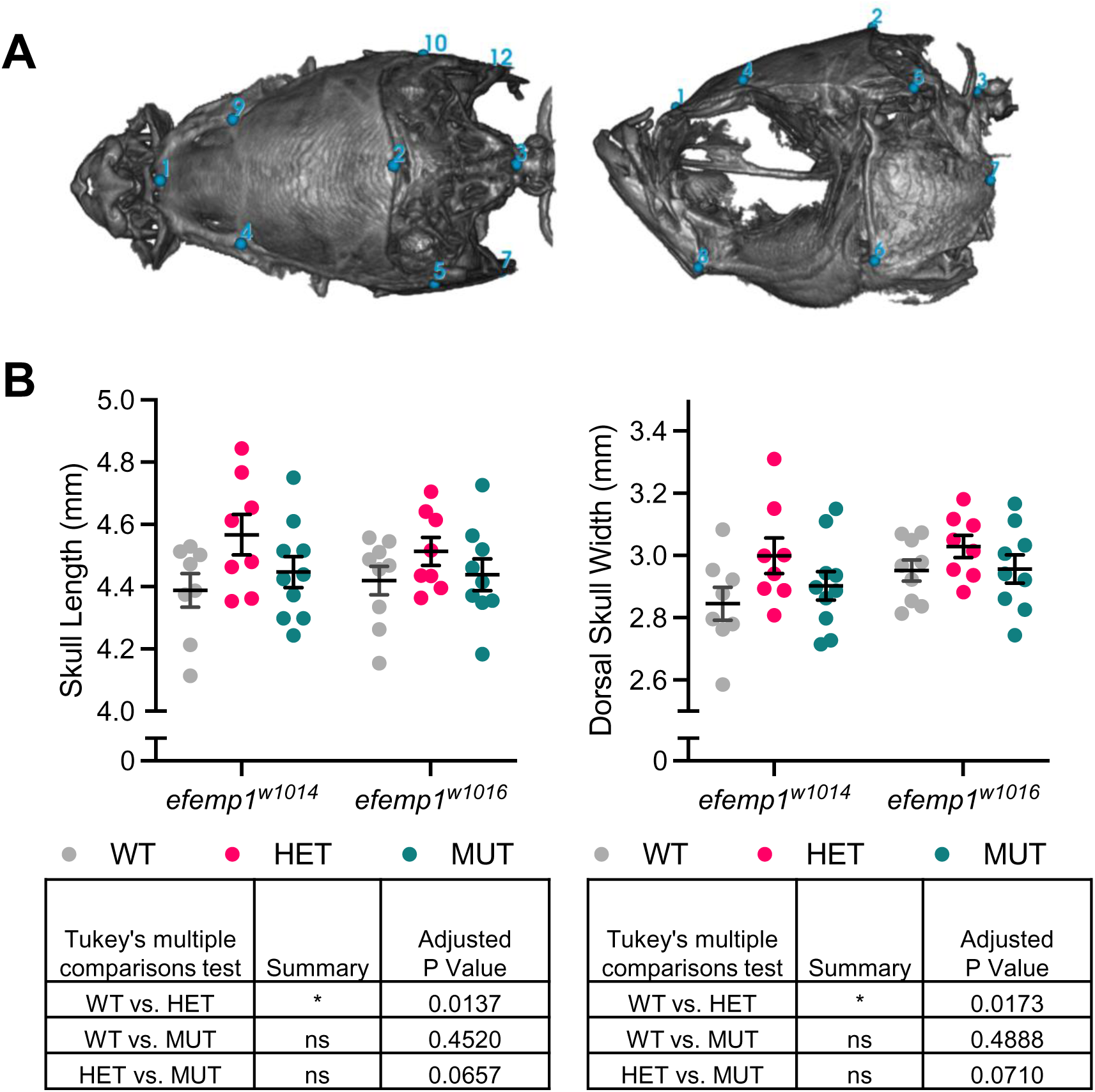
Craniofacial characteristics of zebrafish carrying *efemp1* mutations. **(A)** microCT 3D renderings of a WT fish skull are displayed showing the side and top view. Landmarks 1 – 13 from Table 1 are marked on the rendering. **(B)** We identified significant changes in dorsal posterior skull width and skull length between WT and HET *efemp1^w1014^* and *efemp1^w1016^* fish. The results of Tukey’s multiple comparisons test are shown below the graphs. Error bars represent mean and standard error of the mean. Data points represents a single fish. n = 8-10 fish per group.

## Discussion

In this study, we isolated two new zebrafish *efemp1* mutant alleles and characterized the consequences of heterozygous and homozygous nonsense mutations on vertebral and craniofacial characteristics. By identifying how monoallelic and biallelic variants in *EFEMP1* lead to abnormal skeletal phenotypes, this study helps to better understand the role of EFEMP1 in regulating the skeleton, and the pathophysiological mechanisms underlying the spectrum of genotype-phenotype relationships seen in *EFEMP1*-associated HDCT’s.

While this study was being conducted, an *efemp1* zebrafish mutant (*ulg074*) carrying a nonsense mutation was described (Raman et al. 2024). While the study of Raman et al. primarily focused on the function of *efemp1* in cartilage and its potential role in osteoarthritis risk, it is worthwhile to compare how phenotypes for *efemp1^ulg074^* compared to *efemp1^w1014^* and *efemp1^w1016^* examined in this study. The *efemp1^ulg074/ulg074^* zebrafish exhibited a shorter distance between chondrocranial measurements in the anterior in mid-craniofacial region in 5 dpf animals but these measurements were similar to wildtype fish by 10 dpf (Raman et al. 2024). As late adults (1.5 years), *efemp1^ulg074/ulg074^* mutants exhibited a modest, but significant, increase in vertebral tissue mineral density and centrum tissue mineral density. The most striking feature in these animals was a reduced intervertebral disk space. In our analyses, we only had comparable measurements for centrum tissue mineral density, which was not significantly different in our mutants, but the measurement did appear to be trending higher, similar to what was reported in the *efemp1^ulg074/ulg074^* animals. In our *efemp1^w1014^*and *efemp1^w1016^* mutants, we did not observe an obvious reduction in intervertebral disk space. Differences in the skeletal phenotypes might be explained by complete loss of protein in the knockout model while our model is potentially expressing truncated Efemp1 protein. Differences might also be due to age (1.5 years in the study of Raman et al. vs. 3 months in our study). Indeed, similar intervertebral disc phenotypes are known to progressively manifest in aged zebrafish spines (Kague et al. 2021). Finally, in 2 dpf fish, Raman et al. reported specific *efemp1* expression in the brain, pharyngeal area, and notochord (Raman et al. 2024). This in contrast to our findings where *efemp1* was broadly expressed in adult tissues, similar to studies in human where expression of *EFEMP1* has been identified in skeletal muscle, brain, heart, lung, cartilage, and eye (Giltay et al. 1999; Hasegawa et al. 2017). It is possible that *efemp1* expression in zebrafish, while having more specific expression to cartilage or cartilage-like structures in development, and becomes more broadly expressed in adulthood.

Our findings support the potential for loss-of-function in *EFEMP1* to contribute to altered skeletal characteristics. We found that *efemp1* heterozygous and homozygous mutant adult fish are significantly longer than controls, which correlates with individuals carrying *EFEMP1* nonsense variants, who are tall in stature (Bizzari et al. 2020; Driver et al. 2020). From the microCT analysis, we identified significant changes in vertebral microarchitecture in *efemp1* heterozygous and homozygous mutant animals. Our studies suggest the increased body size and centrum length phenotyped we observed in our *efemp1* mutants did not arise during early spine development. Specifically, in juvenile zebrafish, *efemp1* mutants did not have statistically different body size or abnormal mineralization of developing vertebra when compared to control fish. Notably, clinical phenotypes manifest in children with *EFEMP1* variants and hernias appear in adolescent *efemp1^-/-^* mice (McLaughlin et al. 2007; Bizzari et al. 2020; Driver et al. 2020; Verlee et al. 2021). These data suggest that multiple pathophysiological processes, some which initiate in early development, could act in parallel to render the full spectrum of clinical features seen in *EFEMP1*-associated HDCTs.

Similarly, our findings also support the potential for loss-of-function in *EFEMP1* to contribute to altered craniofacial characteristics. With regard to craniofacial measurements, mutants displayed significantly longer skull length and dorsal skull widths. Craniofacial phenotypes described in individuals with *EFEMP1* variants include long face, increased distance between the inner eyelid corners, and down slanted eyes (Bizzari et al. 2020; Driver et al. 2020; Verlee et al. 2021). As described earlier, Raman et al. reported that *efemp1* mutant zebrafish exhibited disrupted craniofacial morphology as larvae. Thus, it is conceivable that the altered craniofacial morphology in adult mutants originates in early development, owing to its role in cartilage. For instance, there is evidence that EFEMP1 binds with TIMP-3 and ECM-1 (Klenotic et al. 2004; Sercu et al. 2009). ECM1 has previously been identified to play a role in endochondral bone formation, which could be disrupted by variants in *EFEMP*1 in humans and murine models (Deckers et al. 2001; Kong et al. 2016). Compared to humans, in zebrafish limited skeletal structures undergo endochondral ossification, whereas a majority of the human skeleton undergoes this process, which could explain some of the differences in *efemp1* phenotypes between fish and humans. One location where endochondral ossification occurs in both humans and zebrafish are in the skull and pharyngeal skeleton in the cranium, respectively, which could be why we see changes in craniofacial morphology in both human and zebrafish (Le Pabic et al. 2022). Work understanding the potential perturbation of EFEMP1 and ECM1 interaction could broaden our understanding of the skeletal phenotypes in our models and in individuals with *EFEMP1* variants.

We found that phenotypic severity was comparable in heterozygous and homozygous mutants, lending support to the notion that heterozygous nonsense variants play a role in the manifestation of clinical phenotypes. Forghani et al. found evidence that *EFEMP1* gene expression was reduced in the affected individuals due to nonsense-mediated mRNA decay (NMD), suggesting that the Marfan-like HDCT was caused by *EFEMP1* haploinsufficiency. In our work, RT-PCR results showed expression of *efemp1* RNA in fish carrying heterozygous and homozygous mutant alleles were similar to that of wildtype fish, suggesting that transcripts for both alleles escaped NMD. As described above, based on the predicted loss of the fibulin-type module in both mutants, it is reasonable to assume that both mutant alleles were loss-of-function alleles (either hypomorph or null). This would support the notion that mutant phenotypes were due to *efemp1* haploinsufficiency. However, we cannot rule out that the truncated N-terminal peptides encoded by selected nonsense alleles have dominant-negative activity, allowing effective neutralization of function of the WT protein (Judge et al. 2004). In this context, much of the literature for EFEMP1 focuses on its role in the retina, as hetero or homozygosity for an *EFEMP1* missense variant (p.R345W) has been found in individuals with Malattia Leventinese (ML)/Doyne honeycomb retinal dystrophy, which is associated with drusen deposition in the human eye (Stone et al. 1999; Hulleman 2016). Future work examining whether overexpression of truncated Efemp1 in a WT background recapitulates MUT phenotypes may help to delineate these possibilities.

This study raises questions related to cellular mechanism by which mutations in *efemp1* alter body size and skeletal characteristics. Prior studies have identified expression of *Efemp1* in murine tissues including developing heart, condensing mesenchymal cells, condensing cartilage and craniofacial bones, and developing vertebra (Ehlermann et al. 2003). Our work identified *efemp1* expression to vary depending on adult tissue type: *efemp1* was well expressed in zebrafish skin, swim bladder, heart, eyes, and testes. In contrast, there was less expression in bone, arguing for a non-cell autonomous mechanism of action. However, it is important to note while it may be expected that tissues expressing the gene of interest at higher levels would be the most severely affected, the correlation between gene expression and manifestation of clinical phenotypes is not always direct (Feiglin et al. 2017). In zebrafish our models, *efemp1* mutations could be disrupting interactions with proteins that are important for bone specific downstream function, whereas the mutations do not disrupt interactions with key proteins in other tissues. In the murine *Efemp1^-/-^*model, phenotypes included accelerated aging phenotypes including advanced bone age (McLaughlin et al. 2007). In this context, it is worthwhile to note that standard length, which was elevated in *efemp1* mutants, is closely correlated with developmental progress in zebrafish (Parichy et al. 2009). Moreover, intervertebral disc degeneration-like phenotypes, which was seen in the 1.5 year old *efemp1* mutants reported by Raman et al, are a phenotypic feature of aged zebrafish. Thus, an interesting avenue for future research would be examining whether hallmarks of aging, such as cellular senescence, are increased in *efemp1* mutants.

Some limitations of our study should be considered. First, it is noteworthy that knockout mice and individuals with nonsense and missense mutations present with hernias as a consistent phenotype (McLaughlin et al. 2007; Bizzari et al. 2020; Driver et al. 2020; Verlee et al. 2021). We did not notice obvious presentation of hernias in *efemp1* mutants in gross examinations. Similarly, in their recently described knockout zebrafish line, hernias were not reported by Raman et al.. These studies highlight the possibility that loss-of-function in *efemp1* does not disrupt connective tissue in zebrafish in the same way as in humans and mice (Raman et al. 2024). Second, there are limitations to understanding the protein expression of *efemp1* in our model, in that antibodies available for probing *efemp1* in zebrafish lysates are not readily available. Third, while part of our goal was to broaden our understanding of *efemp1*’s role in relationship to the human condition, without comparable knockin models of patient variants or skeletal phenotype measurements in humans, we are limited in our interpretation of a direct relationship to the Marfan-like condition. Thus, animal models harboring variants observed in individuals with *EFEMP1*-associated HDCTs are warranted.

## Acknowledgements

AEG is supported by the Biological Mechanisms for Healthy Aging Training Grant NIH/NIA T32 AG066574. Research reported in this publication was supported by National Institute of Arthritis and Musculoskeletal and Skin Diseases (NIAMS) under Award Numbers AR074417 and the Office Of The Director (OD) of the National Institutes of Health to RYK.

## Notes

### Competing Interest Statement

The authors have declared no competing interest.

