## Supplementary table and figures for "Dominant nonsense mutations in efemp1 alter vertebral and craniofacial characteristics in adult zebrafish"

| | Genotype:allele interaction p-val | Genotype p-val | Allele p-val |
|---|---|---|---|
| Juvenile Standard Length | 0.3841 | 0.8104 | 0.5401 |
| Adult Standard Length | 0.9063 | 0.0128 | 0.2381 |
| Juvenile Calcein Measurements | 0.6838 | 0.5671 | 0.5396 |
| FishCuT Centrum Volume | 0.5481 | 0.0182 | 0.1705 |
| FishCuT Haemal Arch Volume | 0.7327 | 0.1046 | 0.7143 |
| FishCuT Neural Arch Volume | 0.6318 | 0.1021 | 0.7515 |
| FishCuT Centrum TMD | 0.9079 | 0.134 | 0.1994 |
| FishCuT Haemal Arch TMD | 0.9467 | 0.1026 | 0.26 |
| FishCuT Neural Arch TMD | 0.8901 | 0.3438 | 0.2873 |
| FishCuT Centrum Th | 0.1525 | 0.0987 | 0.8506 |
| FishCuT Haemal Arch Th | 0.1028 | 0.3065 | 0.011 |
| FishCuT Neural Arch Th | 0.6763 | 0.3148 | 0.7372 |
| FishCuT Centrum Length | 0.8158 | 0.0238 | 0.7965 |
| Craniofacial Skull Width | 0.1331 | 0.0373 | 0.0036 |
| Craniofacial Frontal/Parietal Bone Length | 0.8573 | 0.489 | 0.0034 |
| Craniofacial Dorsal Skull Width | 0.7014 | 0.0477 | 0.0952 |
| Craniofacial Ventral Skull Width | 0.1365 | 0.0601 | 0.0025 |
| Craniofacial Distance Between Anguloarticulars | 0.0625 | 0.1474 | 0.0003 |
| Craniofacial Epiphyseal Bar Length | 0.5261 | 0.6314 | 0.6166 |
| Craniofacial Skull Length | 0.7283 | 0.039 | 0.8112 |

**Supplementary Table 1. P-values of 2-way ANOVA.** Results of interaction, genotype, and allele p-values for each measurement in this study. All measurements were found to have no significant genotype:allele interaction, suggesting that the effects of *efemp1* mutations for both alleles were similar. Measurements that had a significant p-value for genotype are displayed as main text figures. Other measurements can be found in the supplementary figures.

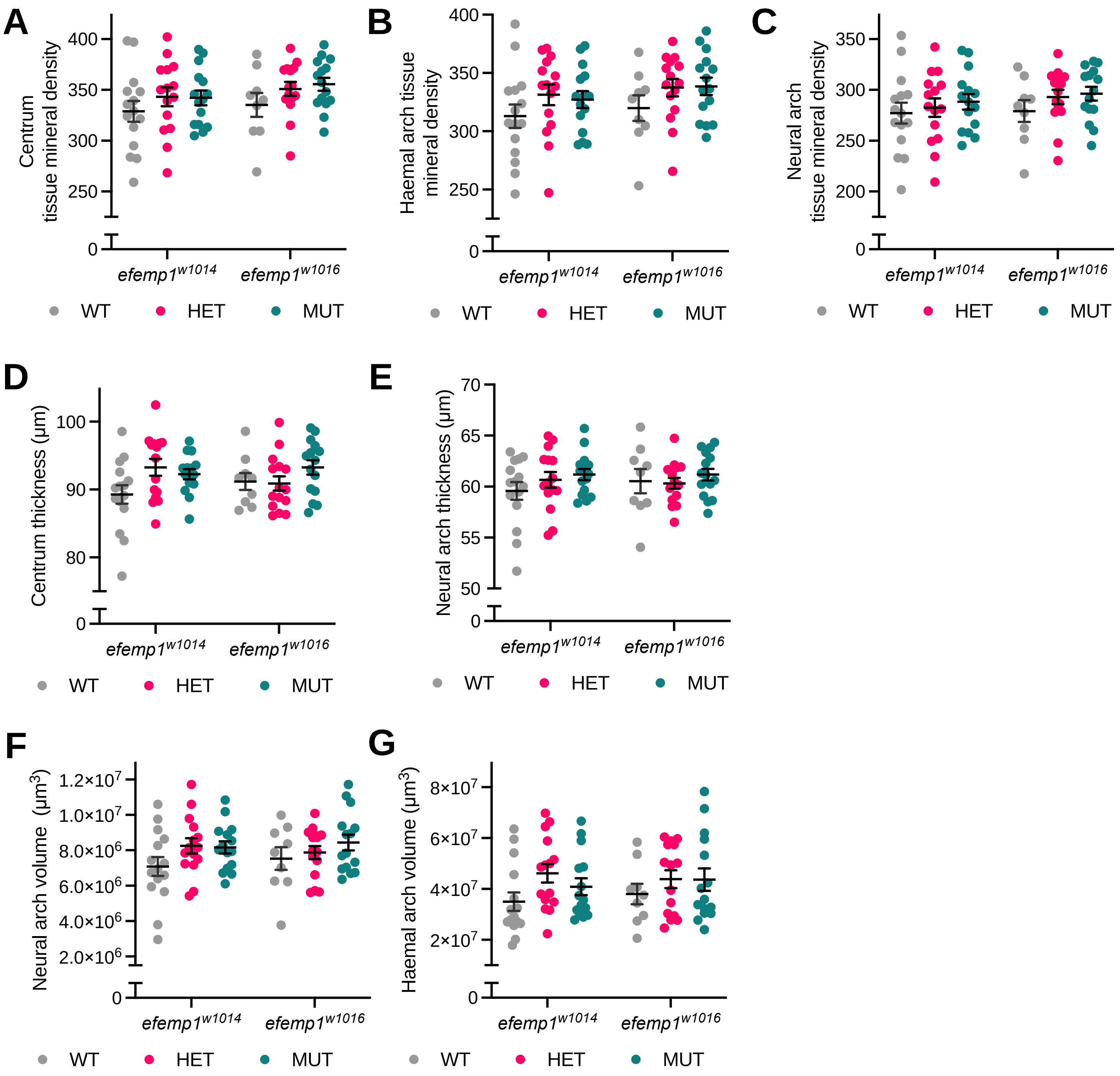


**Supplementary Figure 1. Vertebral measurements of adult zebrafish carrying *efemp1* mutations.** FishCuT analysis results for (A) centrum tissue mineral density, (B) haemal arch tissue mineral density, (C) neural arch tissue mineral density, (D) centrum thickness, (E) neural arch thickness, (F) neural arch volume, and (G) haemal arch volume in *efemp1*$^{w1014}$ and *efemp1*$^{w1016}$ fish. Error bars represent mean and standard error of the mean. Data points represents the average measurement of a single fish for 16 vertebra. n = 9-15 fish per group.

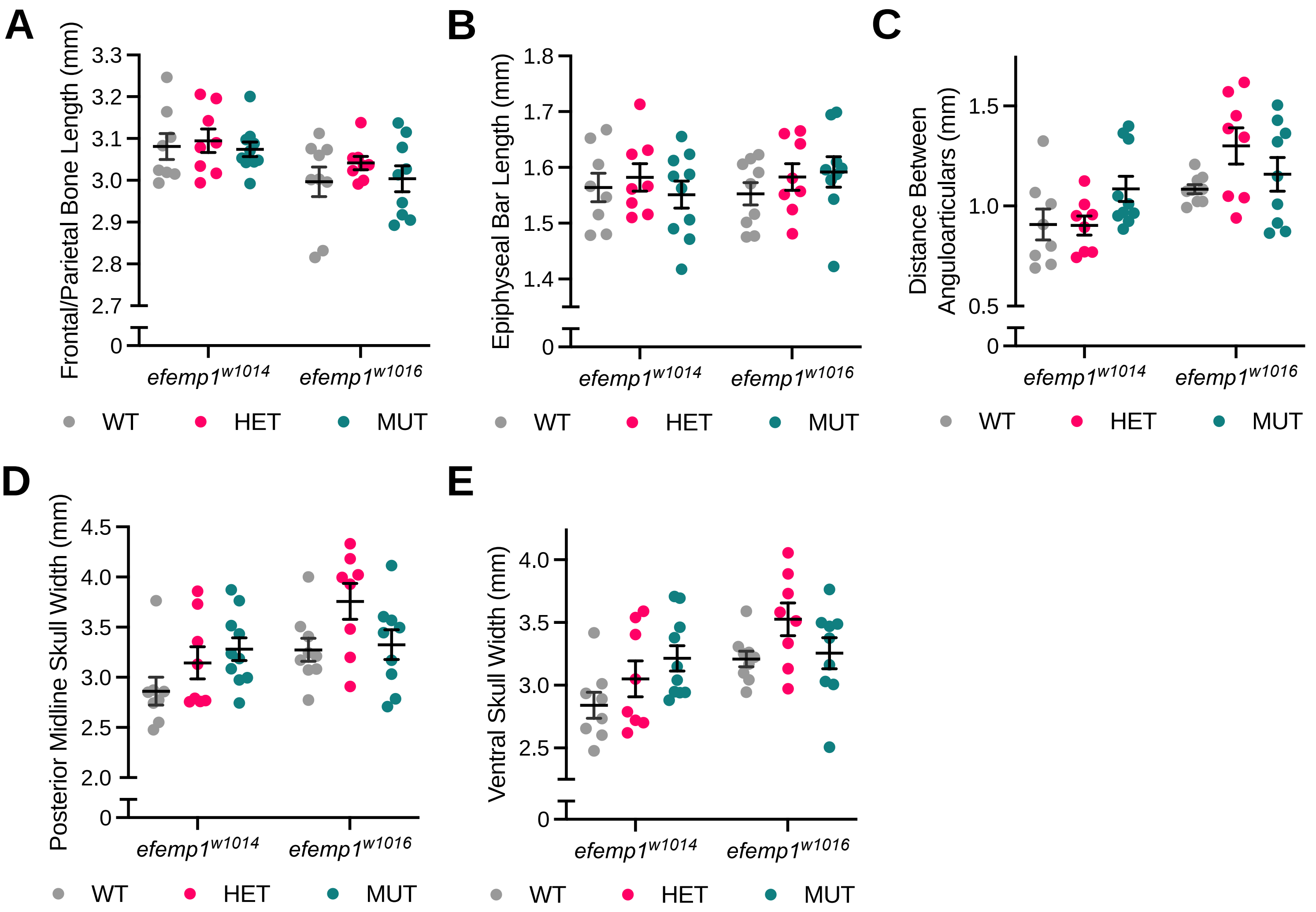


**Supplementary Figure 2. Craniofacial measurements of zebrafish carrying *efemp1* mutations.** Craniofacial measurements for (A) Frontal/Parietal Bone Length, (B) Epiphyseal Bar Length, (C) Distance Between Anguloarticulars, (D) Posterior Midline Skull Width, and (E) Ventral Skull Width. Anatomical points used for each measurement are listed in tables 1 and 2. Error bars represent mean and standard error of the mean. Data points represents a single fish. n = 8-10 fish per group.
